# In vitro immunization approach to generate specific murine monoclonal IgG antibodies

**DOI:** 10.1101/2020.11.24.393728

**Authors:** Sophia Michelchen, Burkhard Micheel, Katja Hanack

**Author notes:** E-mail addresses.

## Abstract

Generating monoclonal antibodies to date is a time intense process requiring immunization of laboratory animals. The transfer of the humoral immune response into *in vitro* settings shortens this process and circumvents the necessity of animal immunization. However, orchestrating the complex interplay of immune cells *in vitro* is very challenging. We aimed for a simplified approach focusing on the protagonist of antibody production: the B lymphocyte. We activated purified murine B lymphocytes *in vitro* with combinations of antigen and stimuli. Within ten days of culture we induced specific IgM and IgG antibody responses against a viral coat protein. Permanently antibody-producing hybridomas were generated. Furthermore we used this method to induce a specific antibody response against *Legionella pneumophila*. We thus established an effective protocol to generate monoclonal antibodies *in vitro*. By overcoming the necessity of *in vivo* immunization it may be the first step towards a universal strategy to generate antibodies from various species.

## 1. Introduction

Antibodies are important tools in biotechnology and medicine with various applications in research, diagnosis and also increasingly in therapeutics (Geskin, 2015). In the organism a complex interplay of dendritic cells (DCs), T lymphocytes and B lymphocytes results in a polyclonal antibody response upon pathogen encounter. Whereas *in vivo* the resulting heterogenous mixture of antibodies in the serum is crucial for the clearance of an immunogen, the biotechnological usage often requires monoclonal antibodies with preferably high affinity and specificity for an antigen of choice. Generating such an antibody to date is a time and labor intensive process.

The hybridoma technique as one of the most widely used methods for antibody production utilizes the *in vivo* immune reaction of a laboratory animal upon immunization. The animal is challenged several times with the antigen of choice. This immunization comprises several weeks and also requires a considerable amount of antigen. Once a satisfying serum titer is reached the animal’s spleen cells are fused with myeloma cells to generate stable hybridoma cell lines that secrete antibodies specific for the administered antigen. Immunization of the living organism and being dependent on spleen cells obviously limits this method to certain species with regards to ethical issues and feasibility. Furthermore, immunizing with highly immunogenic antigens may even cause premature death in laboratory animals.

Therefore, the immune reaction taking place *in vivo* does not leave any opportunity to intervene and underlies a multiplicity of biological factors. To circumvent and reduce these disadvantages a transfer to *in vitro* settings would be favourable. The transfer of antigen-specific immune responses to *in vitro* conditions has first been proposed by Mishell and Dutton in 1966 who induced an antibody response of unimmunized spleen cells. In 1988 Borrebaeck et al. published a general method to produce monoclonal antibodies from human lymphocytes *in vitro*. There have since been several approaches and protocols to induce antigen specific immune responses *in vitro* with varying success and reproducibility. Successful attempts as shown by Wand et al. (2011), Kato et al. (2012) and Inagaki et al. (2013) implied using splenocytes or a combination of DC, T and B lymphocytes leading to the production of specific antibodies *in vitro*. This method allows monitoring of the process while minimizing the time and amount of antigen necessary for specific antibody production. It therefore offers several advantages compared to the common *in vivo* immunization. However, this procedure still involves a number of different cell types that have to be specifically activated and thus factors influencing the outcome in *in vitro* immunization.

To simplify this approach we established a method excluding immune cells other than the cells needed for antibody production, thereby minimizing the number of factors involved and creating a controllable environment for the *in vitro* immune response. In the present study, we describe the establishment of this simplified approach in which only B lymphocytes were specifically activated *in vitro*. In the organism B lymphocytes may be activated by the antigen only (T cell independent) or by a combination of signals from specifically activated T cells and antigen (T cell dependent). T cell independent activation of B lymphocytes mostly results in the production of low affinity immunoglobulin (Ig) M antibodies. To evoke the production of specific IgG antibodies, T cells are needed to provide activation signals to the antigen experienced B cell, which comprise the secretion of different cytokines and the expression of the CD40-ligand on the T cell surface (Crotty 2015; Gardell and Parker 2017). In accordance with this *in vivo* mechanism, we attempt to mimic these activation signals in an *in vitro* setting to efficiently activate B lymphocytes and induce a specific antibody response. Therefore B cells were isolated from naïve mice and cultured with supplements mimicking the *in vivo* conditions of antigen-specific activation. B lymphocyte activation was shown in proliferation assays and by monitoring surface marker expression. Specific IgM responses in *in vitro* cultures were detected from day 3 on, specific IgGs from day 9 onwards. By fusion of antigen-specific B lymphocytes with myeloma cells we were able to generate stable hybridoma cell lines secreting antigen-specific IgM and IgG antibodies. To test if the activation protocol is applicable with different types of antigens we challenged murine B cells with heat-inactivated *Legionella pneumophila* and successfully induced a specific IgG response in our *in vitro* approach.

## 2. Results

### 2.1 Characterization and purity of isolated B lymphocytes

First spleen cells were isolated from naive mice. To eliminate other immune or spleen cells such as T lymphocytes, B lymphocytes were purified through a negative selection with antibody-coated magnetic beads. Purity of B lymphocytes was verified by flow cytometric analysis upon staining CD19, CD4 and CD11c surface markers. All cells were found positive for CD19, but negative for CD4 and CD11c, confirming a pure B lymphocyte population (Fig. 1).

**Fig. 1:**
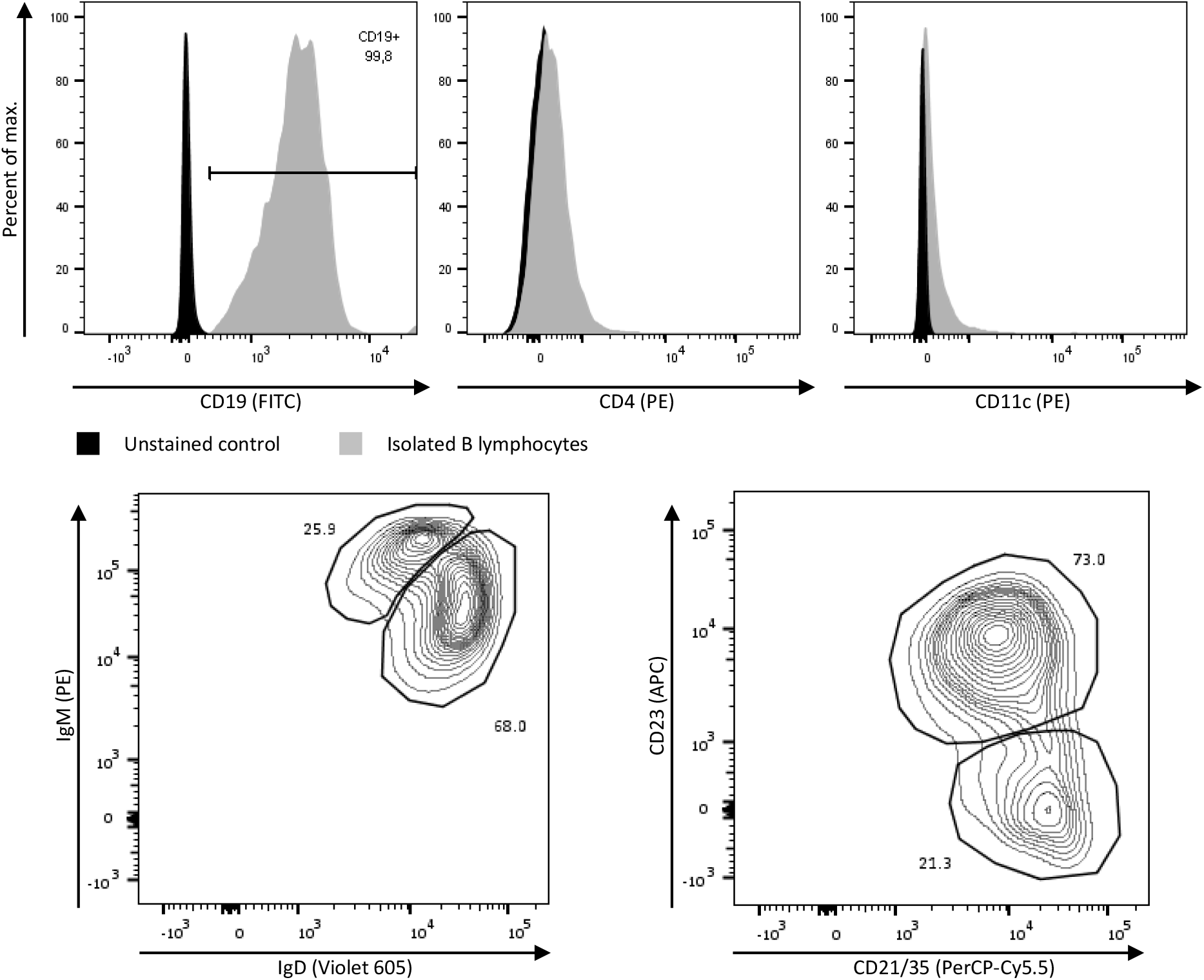
B lymphocytes were isolated from spleen cells of naïve mice by magnetic cell separation. Cells were stained with fluorescein isothiocyanate-labelled anti-CD19, phycoerythrin-labelled anti-CD4, -CD11c and -IgM, Violet 605-labelled anti-IgD, allophycocyanin-labelled anti-CD23 and peridinin chlorophyll-cyanine 5.5-labelled anti-CD21/35 antibodies. By flow cytometric analysis the isolated cells were found to be B lymphocytes only (CD19+) of marginal zone (CD19^+^ IgM^+^ IgD^+^ CD23^−^ CD21^+^) and follicular B lymphocyte (CD19^+^ IgM^+^ IgD^+^ CD23^high^ CD21^+^) phenotype.

Furthermore we investigated the phenotype of the isolated splenic B cells by analyzing the surface expression of IgD, IgM, CD21/CD35, CD23 in combination with CD19. Flow cytometric analysis revealed that all cells expressed IgM, IgD and CD21/35 on their surface, but there were two populations to distinguish. IgM^high^ IgD^+^ cells were further found to be CD23^−^, whereas IgD^high^ IgM^+^ cells were shown to be CD23^+^, indicating the phenotypes of marginal zone (CD19^+^ IgM^+^ IgD^+^ CD23^−^ CD21^+^) and follicular B lymphocytes (CD19^+^ IgM^+^ IgD^+^ CD23^high^ CD21^+^).

### 2.2 Proliferation of B lymphocytes during *in vitro* stimulation

To induce *in vitro* B lymphocyte activation, cells were stimulated with a combination of VP1, anti CD40-antibody, LPS and IL7 for five days. A subsequent restimulation with the cocktail of stimulants for another five days followed and IL4 was added. To investigate whether B lymphocytes proliferate upon stimulation, isolated B lymphocytes were labelled with CFSE and flow cytometric analysis was performed at different time points. More than 90% of the cells stimulated with antigen, anti CD40-antibody, IL4 and LPS or IL7 proliferated upon stimulation. By day 7 the majority of stimulated B lymphocytes had divided at least three times as shown in Fig. 2. Proliferation however seemed to decline after day 7 as there is no considerable difference seen between proliferation status of cultures from day 7 and day 9 (not shown). Number of living cells did not increase beyond input numbers but stayed roughly constant throughout the stimulation. Unstimulated control cells did not show any proliferation. Also stimulation with antigen only had no effect on B cell growth. In fact these cultures constantly decreased in numbers of living cells and cells had eventually died out by day 9.

**Fig. 2:**
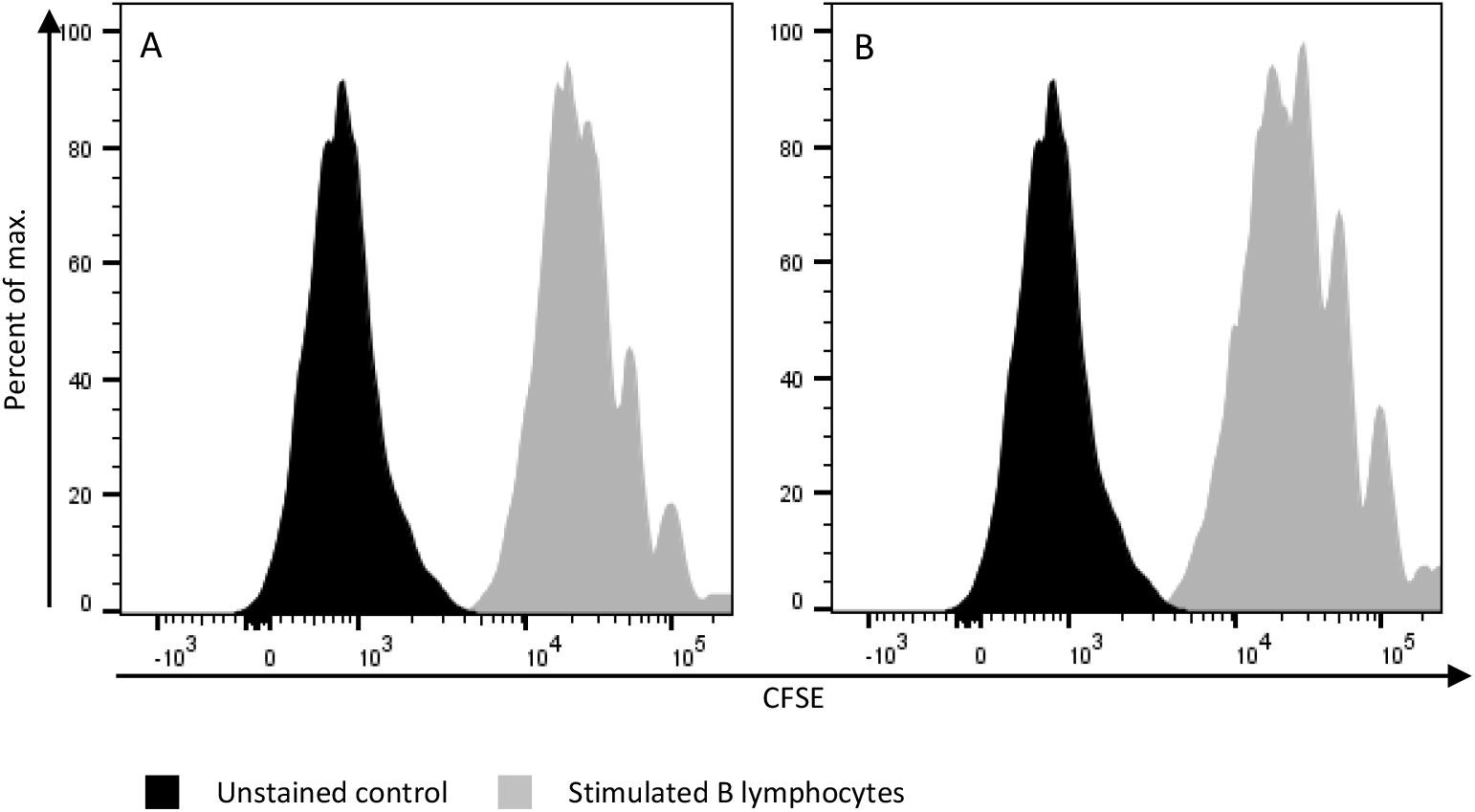
Isolated B lymphocytes were stained with CFSE and stimulated *in vitro* with 10 μg/ml VP1 and 1 μg/ml LPS (A) or 2 ng/ml recombinant murine IL7 (B) for ten days. Proliferation as indicated by the decreasing fluorescent intensity of CFSE is shown here on day 7 of *in vitro* stimulation. In both samples the majority of cells has divided at least three times and a total of four to five daughter generations can be traced.

### 2.3 Activation of B lymphocytes

Upon activation of B lymphocytes the surface marker expression changes. We therefore examined the stimulated cells for their expression patterns throughout the stimulation. Surface markers CD69, CD80, CD86, CD138 and surface IgG were labelled and analyzed at different time points. Stimulated cells showed a drift towards higher expression of activation markers CD69, CD80 and CD86 within ten days indicating a successful *in vitro* activation (Fig. 3). A fraction of the isolated cells already expressed surface IgG at an intermediate level initially. After ten days of *in vitro* stimulation the expression had shifted to a considerably higher level in all cells. However, the expression of CD138 as a prominent marker of plasma cells did not or only marginally increase.

**Fig. 3:**
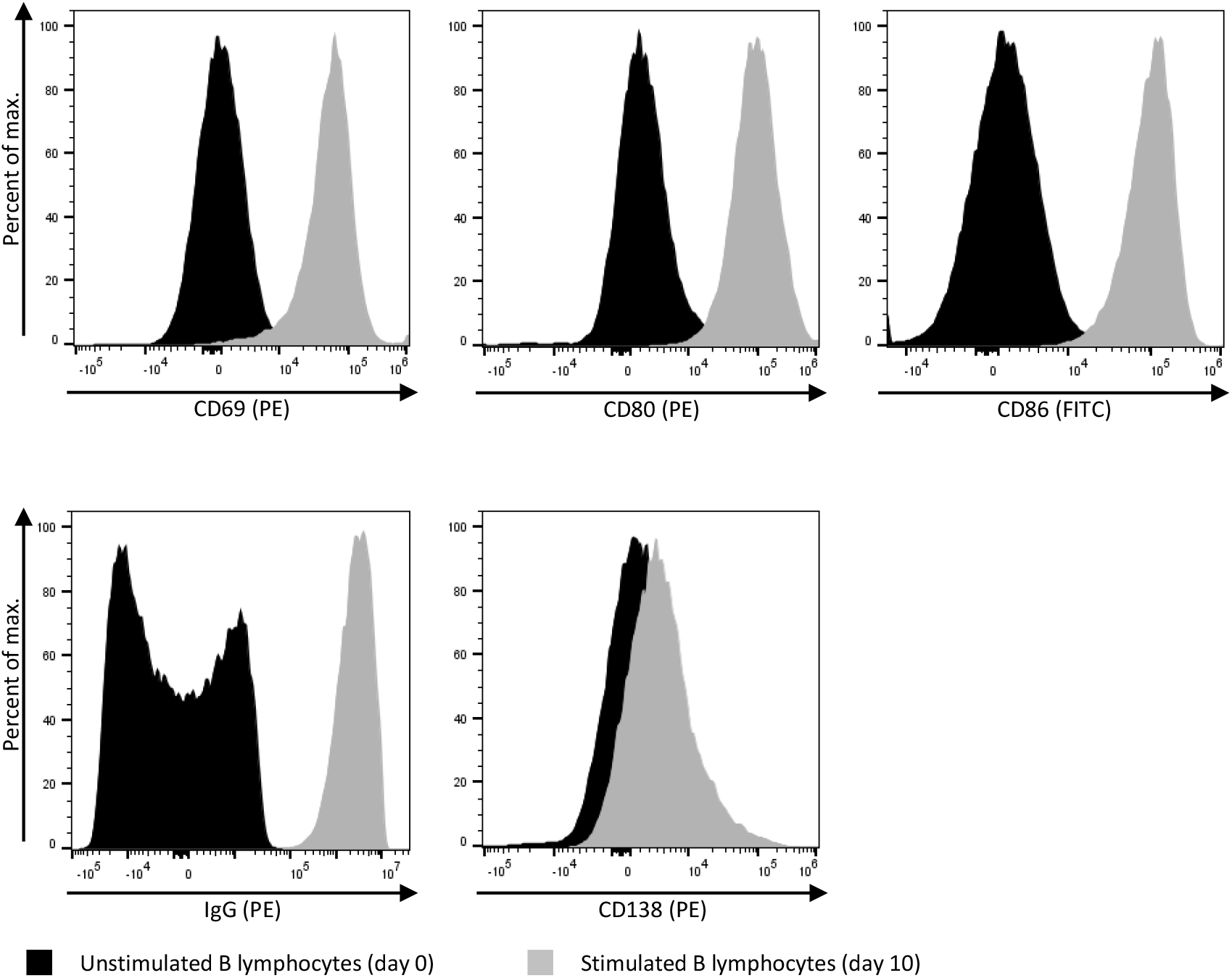
*In vitro* stimulated B lymphocytes were stained for expression of surface activation markers CD69, CD80, CD86, CD138 and surface IgG. Compared to unstimulated cells a strong increase in CD69, CD80, CD86 and surface IgG expression was observed in cultures stimulated with antigen (VP1), anti CD40-antibody, IL4, LPS or IL7. Expression of CD138 did not change remarkably within the ten days of *in vitro* stimulation.

### 2.4 Induction of antigen-specific antibodies by *in vitro* activation

During stimulation, the presence of antibodies specific for the antigen VP1 among the different culture settings was determined daily by ELISA (Fig. 4). Specific IgM antibodies were detected from day 3 on in cultures stimulated with antigen alone or antigen, anti CD40-antibody, IL4, LPS or IL7. Specific IgG secretion only evoked upon secondary antigen stimulation and peaked on day 10 of *in vitro* activation similar to IgM. In stimulated cultures we detected a significantly higher level of VP1-specific IgM and IgG antibodies compared to unstimulated control cultures. While cells stimulated with antigen alone showed highest levels of both specific IgM and IgG these cells suffered from poor viability and did not proliferate. These data show a successful activation of naive B lymphocytes *in vitro* induced by signals mimicking *in vivo* T cell activation.

**Fig. 4:**
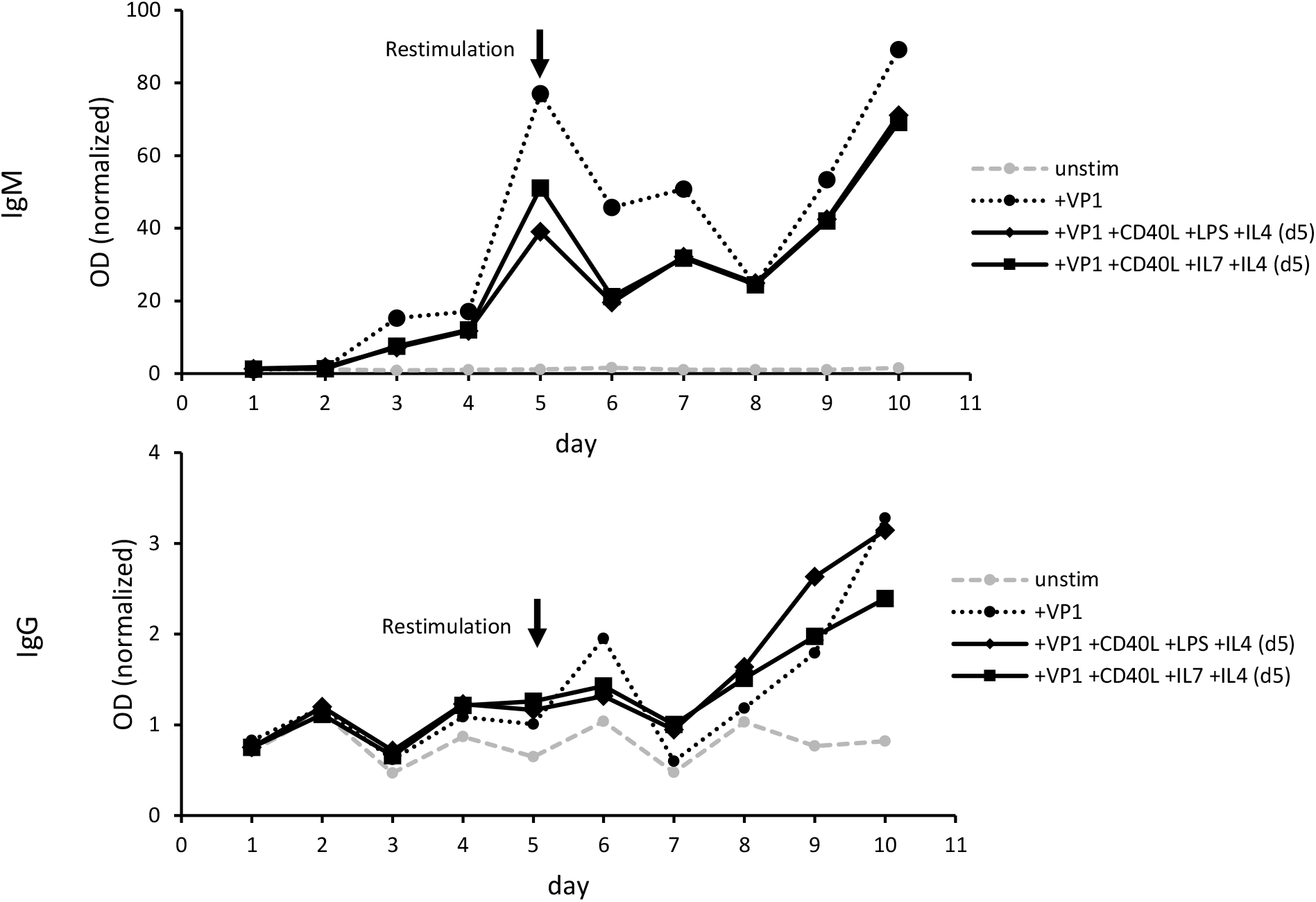
Murine B lymphocytes isolated from naive mice were stimulated *in vitro* with antigen (VP1), anti CD40-antibody, IL4, LPS or IL7 and restimulated after five days. Supernatants were tested daily for specific IgM and IgG antibodies. A strong IgM response arose from day 3 onwards in samples stimulated with antigen (VP1), anti CD40-antibody, IL4, LPS or IL7 and peaked in a signal 90-fold higher than that of unstimulated cells on day 10. Antigen-specific IgG antibodies were detected on day 9 in the same samples.

### 2.5 Establishment of a monoclonal antibody with *in vitro* activation

We then attempted to establish a monoclonal antibody by using our activation method in combination with hybridoma technology. Fusion to generate hybridomas from the *in vitro* activated B lymphocytes was carried out five days after restimulation, i.e. after ten days of *in vitro* activation. Fusion efficiency was very high among cultures stimulated with antigen, anti-CD40 antibody, IL4 and LPS or IL7 and several clones could be established. After the selection process we identified several monoclonal hybridoma cell lines stably producing VP1-specific IgG antibodies of IgG2b subclass (Fig. 6). For all antibodies derived from *in vitro* immunization positive signals (OD > 0.2) were obtained at concentrations of 0.625 μg/ml or higher, however signal intensity is lower compared to an antibody (P157) obtained from conventional *in vivo* immunization with VP1 (Fig. 5).

**Fig. 5:**
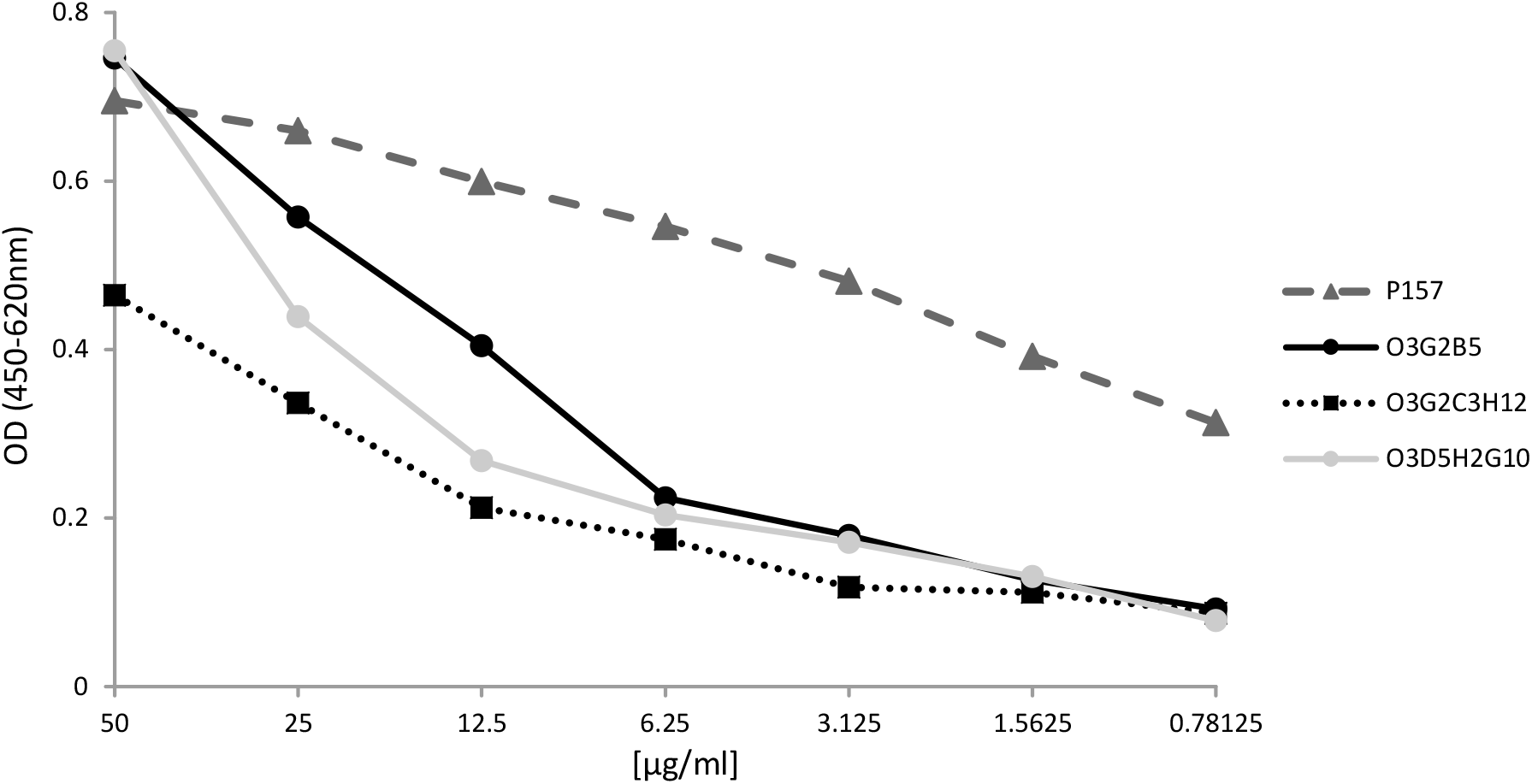
Purified monoclonal antibodies were tested against the antigen VP1. Antibodies O3G2B5, O3D5H2G10 and O3G2C3H12 were obtained from generated hybridoma cell lines after *in vitro* immunization of B lymphocytes with O3G2B5 showing the highest binding ability against the antigen. In comparison, P157 is an antibody derived from conventional *in vivo* immunization with VP1. Antibody concentrations ranged from 50 to 0.78125 μg/ml.

**Fig. 6:**
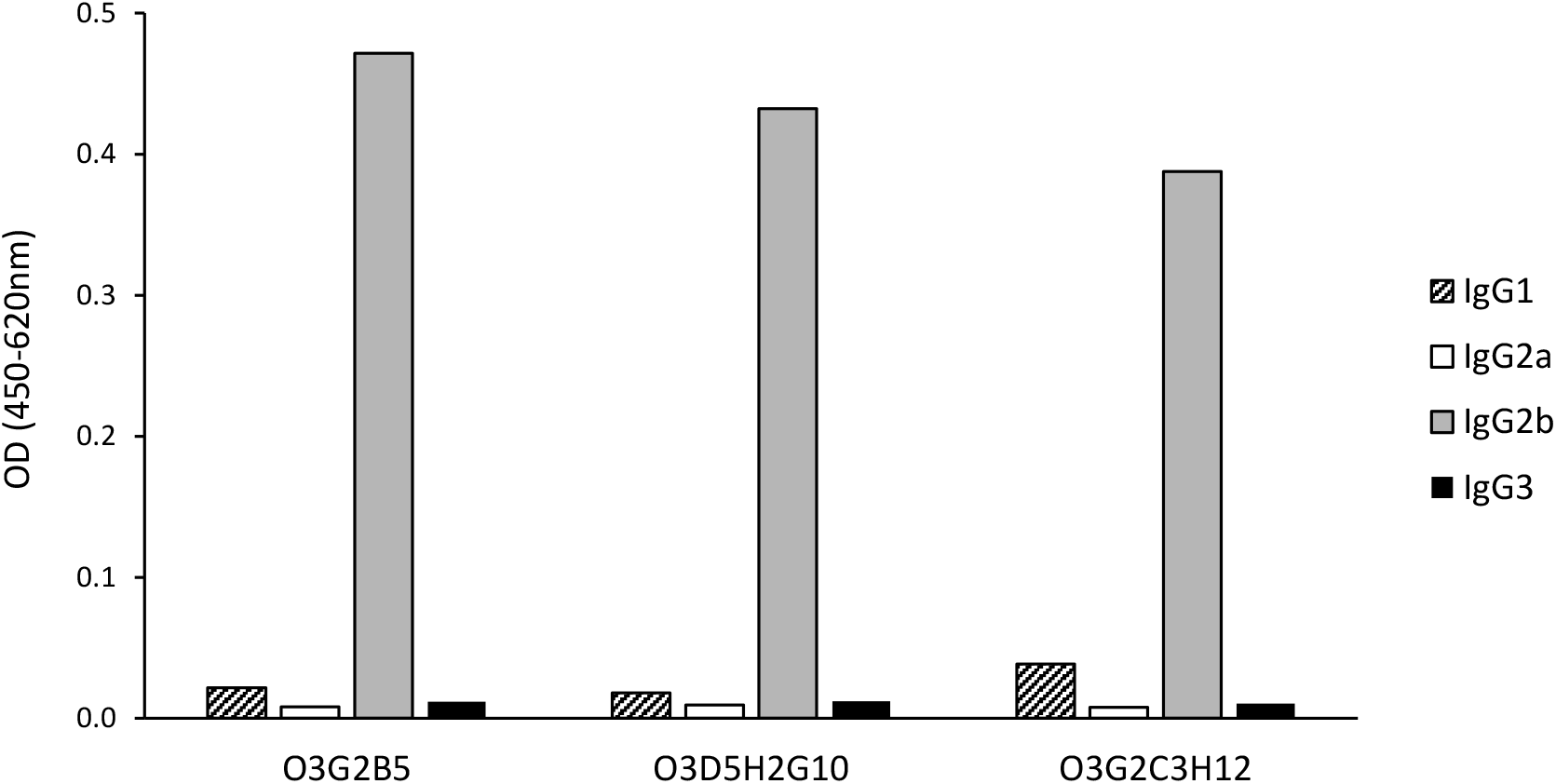
Subclasses of purified monoclonal antibodies were determined by ELISA. Subclasses were detected by subclass specific biotin-labeled antibodies and subsequent streptavidin-conjugated horse radish peroxidase. All antibodies were identified as IgG2b.

### 2.6 Induction of *in vitro* antibody response against Legionella pneumophila

Next we challenged murine B lymphocyte cultures with heat-inactivated *L. pneumophila* to test transferability of the activation protocol to different antigen types. Culture supernatants were tested for the presence of specific IgG antibodies by flow cytometric analysis. Therefore *L. pneumophila* were incubated with the supernatants of stimulated versus unstimulated B lymphocytes and subsequently stained with a PE-labelled anti-mouse IgG secondary antibody. Supernatants from cultures stimulated with both *L. pneumophila*, anti CD40-antibody, IL4 and LPS or IL7 (Fig. 7) clearly contained IgG antibodies capable of binding to and labelling *L. pneumophila* whereas unstimulated B lymphocyte cultures did not produce such.

**Fig. 7:**
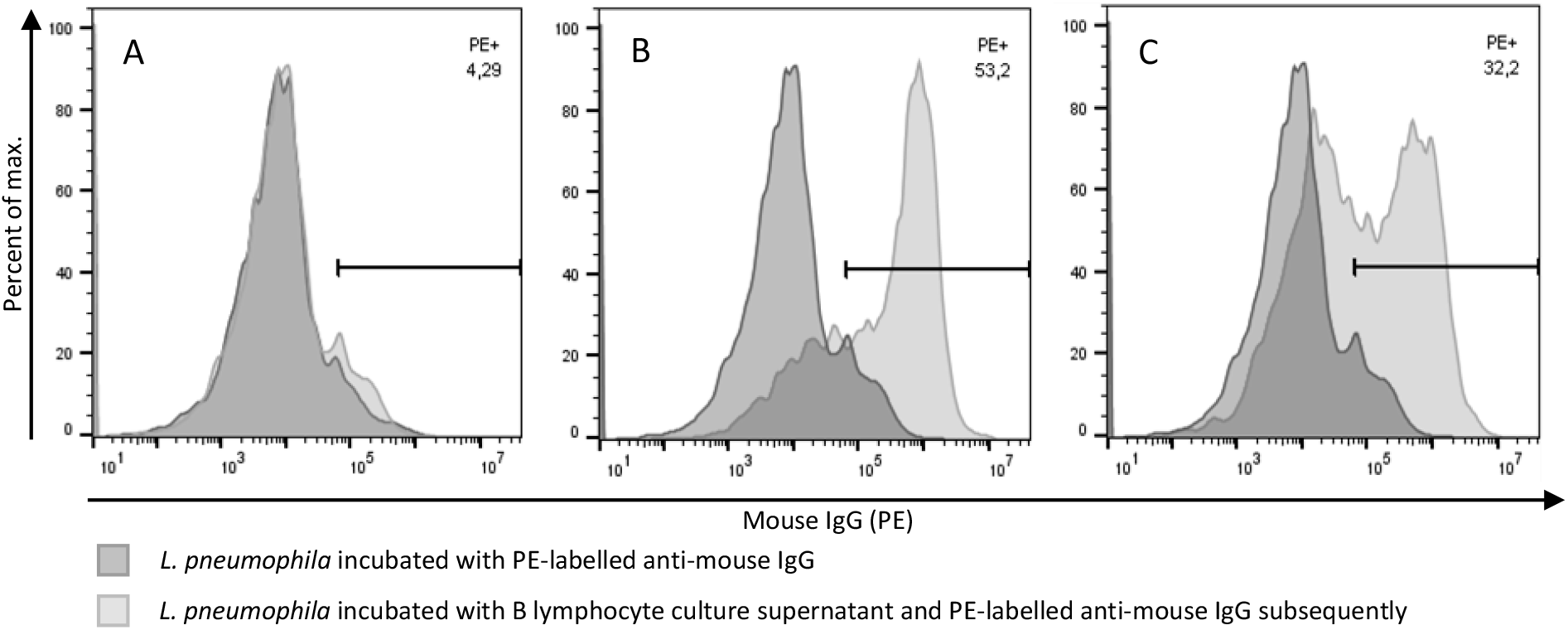
Specific IgG antibodies against *L. pneumophila* by *in vitro* activation of B lymphocytes. Murine B lymphocytes from naive mice were cultured *in vitro* with or without heat-inactivated *L. pneumophila* and a combination of stimuli and restimulated after five days. After ten days of *in vitro* activation supernatants from B lymphocyte cultures were harvested and incubated with heat-inactivated *L. pneumophila*. Upon washing *L. pneumophila* were stained with PE-labelled anti-mouse IgG secondary antibody. A: *L. pneumophila* incubated with supernatant from unstimulated murine naïve B lymphocytes. B and C: *L. pneumophila* incubated with supernatants from B lymphocytes that were stimulated with heat-inactivated *L. pneumophila*, anti CD40-antibody, IL4 and LPS (B) or IL7 (C), respectively. Samples labelled with secondary anti-mouse IgG antibody only are depicted in dark grey; shown in light grey are samples incubated with supernatant and subsequently with secondary antibody.

## 3. Discussion

In this study we showed the *in vitro* induction of a specific IgG antibody response by *in vitro* activation of murine B lymphocytes. For the activation of B lymphocytes *in vivo* a coordinated action of DCs and T lymphocytes is necessary to trigger a specific IgG response by B lymphocytes. *In vivo*, DCs present antigen to T and probably also B lymphocytes. Since B lymphocytes also function as antigen presenting cells, they might take over antigen presentation themselves *in vitro* instead of DCs (Rodríguez-Pinto, 2005). Whether specific subsets of B lymphocytes are responsible for this function is yet unknown and has to be clarified (Chen and Jensen, 2008; den Haan et al., 2014; Hong et al., 2018; Kambayashi and Laufer, 2014). *In vivo*, T lymphocytes take up processed antigen from DCs and present a complex of MHCs and antigenic peptides to B lymphocytes. In combination with a range of further signals, this process activates antigen-specific B lymphocytes to produce antibodies whose subclass depends on these signals. T lymphocytes are also partially responsible for the avoidance of an immune response against the body’s own antigens. As far as we can consider, autoimmune reactions do not play a role during *in vitro* in contrast to *in vivo* immunization, thus allow for a wider spectrum of antibodies.

The main obstacle in *in vitro* immunization is finding a method to activate B lymphocytes to produce antibodies of the wanted subclass and specificity. *In vivo*, B lymphocytes can be activated in a T cell-independent manner by antigen only leading to the production of low affinity IgM antibodies. This was transferred to *in vitro* settings repeatedly (Ait Mebarek et al., 2013; Federspiel et al., 1991; Wohlleben et al., 1996). Those methods however lack an efficient activation of B cells resulting in a limited number of sufficiently matured antibody producing cells, and thus struggle to deliver specific antibodies other than IgM. Other studies succeeded to induce antigen specific IgG responses *in vitro* by immunizing a heterogeneous population of splenocytes by exposure to antigen only or with a combination of cytokines and stimulative antibodies (Ravi et al., 2007; Inagaki et al., 2013), but failed to deliver a reliable and reproducible protocol presumably due to variation in antigen specific and successfully activated immune cell populations. To overcome these difficulties we aimed to reduce the impact of variables. In the present study we therefore tried to specifically activate murine B lymphocytes alone *in vitro*.

Splenic B lymphocytes were isolated from naive C57Bl/6 mice and purity was shown by FACS analysis as no DCs or T cells could be detected in the cell suspension. *In vivo* signals of antigen-specific T cell mediated activation were mimicked *in vitro* by using a range of immunomodulators in different combinations. To circumvent the DC function, B lymphocytes were challenged with a surplus of soluble model antigen VP1. B cell activation *in vitro* was verified by changes in surface receptor expression, proliferation and antibody production. Indeed, the addition of anti CD40-antibody, IL4 and LPS or IL7 to the culture medium in combination with the antigen led to an efficient activation of B lymphocytes *in vitro* and moreover resulted in an increased production of anti-VP1 IgG antibodies by the B lymphocytes. Furthermore we were able to equivalently induce specific IgG responses when activating B lymphocytes *in vitro* in similar settings with other antigens such as *L. pneumophila*. Without T cell-substituting ingredients the specific IgG response was significantly lower as was cell viability. Moreover, it was not possible to sustain living cells throughout the stimulation to generate hybridoma cells from. From B lymphocyte cultures supplemented with antigen and T cell-replacing ingredients we were able to produce a number of stable hybridomas synthesizing VP1 specific antibodies of IgG2b subclass. Hence, it is proven possible to obtain monoclonal antigen-specific antibodies with our *in vitro* method. While antibodies generated in *in vitro* immunization produced positive ELISA signals at low concentrations, overall signal intensities were lower compared to an anti VP1 antibody obtained from conventional *in vivo* immunization. This can be attributed to the shortened time period of immunization and thus incomplete affinity maturation. In contrast to the conventional method of antibody production by *in vivo* immunization the procedure was shortened from several months to a few days while lower amounts of antigen were needed and potential distress for experimental animals was minimized.

Several aspects require further exploration. Since VP1 as a viral protein has adjuvant effects it has to be clarified whether other proteins and haptens can induce an *in vitro* antibody response in the same setting. Additionally, the method is to be tested for the human system which could be a substantial breakthrough. Eventually, this may be a universal procedure to produce specific antibodies *in vitro* minimally invasive from various species.

## 4. Material and methods

### 4.1 Purification and culture of murine splenic B lymphocytes

Spleens were removed from naïve C57Bl/6j mice (8-12 weeks). Single cell suspensions of splenocytes were obtained by dispersion through a 40 μM cell strainer (VWR, Radnor, Pennsylvania, USA). B cells were isolated by negative selection using the B Cell Isolation Kit (Miltenyi Biotec, Bergisch-Gladbach, Germany). The procedure was carried out following the instructions of the manufacturer. Purified B cells were cultured in 24 well plates (TPP, Trasadingen, Switzerland) at 10^6^ cells/well in 1 ml RPMI 1640 complete medium (Thermo Fisher Scientific, Waltham, Massachusetts, USA) supplemented with 10 % heat-inactivated fetal calf serum (FCS, Thermo Fisher Scientific, Waltham, Massachusetts, USA), 2 mM L-glutamine (Roth, Karlsruhe, Germany) and 50 μM 2-mercaptoethanol (Roth, Karlsruhe, Germany) and kept at 37 °C and 5 % CO_2_.

### 4.2 Proliferation assay

Following purification, B lymphocytes were stained with carboxyfluorescein succinimidyl ester (CFSE, Thermo Fisher Scientific, Waltham, Massachusetts, USA) according to the protocol published by Quah et al. (2007). In short, cells were resuspended in PBS containing 5 % FCS. CFSE was added to a final concentration of 1.25 μM and cell suspension was mixed immediately. Cells were left to incubate for 5 min at room temperature (RT) in the dark. Subsequently, cells were washed three times by diluting the suspension with PBS/5 % FCS and centrifugation was performed for 5 min at 300 x g. Cells were resuspended in complete medium and used for *in vitro* stimulation. Before analysis, cells were stained with 1 μM PoPro-1 Iodide (Thermo Fisher Scientific, Waltham, Massachusetts, USA) for 10 min at 4 °C in the dark to discriminate living cells. Cells were examined using the Attune Flow Cytometer (Thermo Fisher Scientific, Waltham, Massachusetts, USA). Data were analysed using the FlowJo software (FlowJo LLC, Ashland, Oregon, USA).

### 4.3 In vitro immunization

Following purification or CFSE staining, naïve B lymphocytes were challenged with 10 μg/ml hamster polyomavirus capsid protein (VP1, as described previously in Lütkecosmann et al., 2019) serving as model antigen or 5 x 10^4^ to 5 x 10^6^ heat-inactivated *Legionella pneumophila* (kindly gifted by sifin diagnostics gmbh, Berlin, Germany) per ml. Different combinations of stimuli were added to the culture, i.e. 2 μg/ml rat anti-mouse CD40 antibody (BioLegend, San Diego, California, USA), 2 ng/ml interleukin 7 (IL7) (Miltenyi Biotec, Bergisch-Gladbach, Germany) and/or 1 μg/ml lipopolysaccharide (LPS) (Merck KGaA, Darmstadt, Germany). Control cultures were left untreated. Culture supernatants were collected for the determination of specific antibodies. On day 5 of cultivation, remaining supernatant was reduced to 500 μl. To restimulate the cells, 1 ml of medium containing VP1 or inactivated *L. pneumophila* and the appropriate stimuli was added to each well. Additionally, interleukin 4 (IL4) (Miltenyi Biotec, Bergisch-Gladbach, Germany) was added to some wells at 10 ng/ml.

### 4.4 Flow cytometric analysis of B lymphocytes

Activation status of B lymphocytes was examined by flow cytometric analysis of several surface activation markers. Therefore, cells were harvested and washed in PBS. Cells were then labelled in 300 μl staining buffer using combinations of different antibodies for 30 min at 4 °C in the dark (anti-mouse CD4, CD69 (BD Bioscience, Franklin Lakes, New Jersey, USA), CD19, IgD (Thermo Fisher Scientific, Waltham, Massachusetts, USA), CD11c, CD80, CD86, CD138 (Miltenyi Biotec, Bergisch-Gladbach, Germany), CD21/CD35, CD23, IgM and IgG (BioLegend, San Diego, California, USA)). Cells were then washed as before and resuspended in 300 μl 1 μM PoPro-1 Iodide to obtain the total number of living cells. Upon incubation for 10 min at 4 °C in the dark, cells were examined using the BD FACSAria III (BD Bioscience, Franklin Lakes, New Jersey, USA). Data were analysed using the FlowJo software.

### 4.5 Detection of antigen-specific antibodies

Supernatants were tested daily in enzyme-linked immunosorbent assay (ELISA) for specific IgM and IgG production. Therefore, 96 well plates were coated with 50 μl/well of 5 μg/ml antigen in PBS at 4 °C overnight. Following blocking with 100 μl PBS containing 5 % neonatal calf serum (NCS) for 60 min at RT, 50 μl culture supernatants were added for 45 min at RT. For the detection of antigen-bound antibodies the wells were incubated with 50 μl of horseradish peroxidase (HRP)-labeled anti-mouse IgM or IgG antibodies (Dianova, Hamburg, Germany) for 45 min at RT. Finally, 50 μl substrate solution containing 0.12 mg/ml tetramethylbenzidine (Roth, Karlsruhe, Germany, in 50 mM NaH_2_PO_4_ with 0.04 % CH_4_N_2_O·H_2_O_2_) were added per well and the reaction was stopped using 1 M sulfuric acid. Absorbance was measured at 450 nm with a reference wavelength of 620 nm using the Multiskan FC Microplate Photometer (Thermo Fisher Scientific, Waltham, Massachusetts, USA).

### 4.6 Cell fusion of murine B lymphocytes and purification of monoclonal antibodies

On day 10 of *in vitro* culture and stimulation, cells were harvested for fusion with murine myeloma cell line SP2/0-Ag14 by electrofusion in the presence of polyethylene glycol (Roth, Karlsruhe, Germany) as described previously (Holzlöhner and Hanack 2017). Fused cells were then seeded on 96 well flat bottom plates onto mouse peritoneal cells as feeder layer in hypoxanthine-aminopterine-thymidine (VWR, Radnor, Pennsylvania, USA) medium for the selection process.

Cultures with positive ELISA signals were seeded in limited dilution and inspected microscopically for monoclonal growth. Monoclonal hybridoma were transferred into T75-flasks (VWR, Radnor, Pennsylvania, USA) and supernatants were collected up to 500 ml. IgG antibodies were then purified as described previously (Holzlöhner and Hanack 2017). In short, supernatants were filtered through a 0.45 μm filter (Sartorius Stedim Biotech GmbH, Goettingen, Germany), mixed with binding buffer (4 M NaCl and 1 M glycine NaOH, pH 8.9) and then loaded onto a protein A column. Bound antibodies were eluted with 0.1 M citrate, pH 3.5, and neutralized with 500 μl Tris-HCl, pH 9.0.

### 4.7 Determination of immunoglobulin subclass and direct ELISA

ELISA was carried out as described before. Briefly, 96 well plates were coated with 50 μl/well of 5 μg/ml antigen in PBS at 4 °C overnight. Following blocking with 100 μl PBS containing 5 % NCS, 50 μl of antibody diluted in PBS/5 % NCS (50 – 0.78 μg/ml) were added for 45 min at RT. To determine subclasses of purified antibodies wells were incubated for 45 min with 50 μl of biotin-labeled anti-mouse IgG1, IgG2a, IgG2b or IgG3 antibodies (Jackson ImmunoResearch Europe Ltd, Cambridgeshire, United Kingdom). Subsequently, 50 μl of streptavidin-HRP (Merck KGaA, Darmstadt, Germany) were added and incubated for 45 min. For comparison of antibodies, wells were incubated with 50 μl of HRP-labeled anti-mouse IgG antibodies (Dianova, Hamburg, Germany) for 45 min at RT. Subsequent steps were carried out as described before.

### 4.8 Detection of antibodies specific for Legionella pneumophila

Heat inactivated *L. pneumophila* were washed with PBS at 6500 x g for 5 min and incubated with 100 μl supernatant from stimulated or unstimulated B lymphocyte cultures. Upon washing, *L. pneumophila* were incubated with PE-labelled anti-mouse IgG antibodies for 30 min at 4 °C in the dark. *L. pneumophila* were then washed as before and examined using the Attune Flow Cytometer (Applied Biosystems, Foster City, California, USA). Flow cytometric data were analysed using the FlowJo software.

## Abbreviation

CFSE: carboxyfluorescein succinimidyl ester
DC: dendritic cell
ELISA: enzyme-linked immunosorbent assay
FCS: fetal calf serum
HRP: horse raddish peroxidase
Ig: immunoglobulin
IL: interleukin
LPS: lipopolysaccharide
NCS: neonatal calf serum
RT: room temperature
VP1: hamster polyomavirus capsid protein

## Acknowledgements

We thank the German Federal Ministry of Education and Research for funding the work (03IPT7030X).

## Competing interests

The corresponding author Katja Hanack is owner and manager of new/era/mabs. The other authors declare no competing financial or non-financial interests.

